# Genotype identification and diversity analysis in Kiwifruit (*Actinidia* spp.) using RAPD markers

**DOI:** 10.1101/2020.10.14.339358

**Authors:** Aarti Kamboj, Pushpa Kharb, Agrim Jhilta, Rakshita Singh

**Affiliations:** Department of Molecular Biology, Biotechnology & Bioinformatics, CCS Haryana Agricultural University, Hisar; Himalayan Forest Research Institute, Conifer Campus, Shimla

## Abstract

Kiwifruit (*Actinidia spp*.) is a significant plantation crop belonging to family Actinidiaceae, having deciduous, dioecious, and scrambling vines with chromosome number 2n=58. Dioecy in kiwifruit forms the basis for several breeding programs. The present study was carried out for diversity analysis in kiwifruit genotypes using RAPD markers. 7 kiwifruit genotypes (2 males viz. Allision & Tomuri and 5 females viz. Hayward, Bruno, Allision, Monty & Abbott) were analysed for molecular polymorphism using 94 RAPD primers, out of which 23 primers amplified the genomic DNA in all the genotypes. RAPD data was analysed using NTSYS-pc software and dendrogram construction was done using UPGMA method. Two separate clusters of male and female genotypes were formed. Similarity matrix indices showed maximum similarity between Tomuri (M) and Allision (M) with a similarity coefficient of 0.719 while Abbott (F) and Allision (M) were found to have least similarity having a similarity coefficient of 0.521. Four RAPD primers amplified unique amplicons in Monty, Hayward, Bruno, Allision (M) and Abbott and two primers amplified unique amplicons in Allision (M) & Tomuri (M) along with the male and female plants of Allision genotype respectively. Therefore, these primers can help in distinguishing the genotypes of kiwifruit and can also be validated as putative markers for the sex identification in kiwifruit.

## INTRODUCTION

Kiwifruit (*Actinidia* spp.) is a member of family Actinidiaceae. The genus *Actinidia* consists of more than 50 species and 75 taxa, mostly originated from Southwest China and extensively distributed in eastern Asia. They are mostly deciduous, dioecious, and scrambling vines (Ferguson, 1991). The basic chromosome number in *Actinidia* is X=29, with diploid number 2n=58 (Yan *et al*., 1997). But there exist ploidy variations among different species, i.e. they may be tetraploid (e.g. *A. chinensis*), hexaploid (e.g. *A. deliciosa*) or octaploid (e.g. *A. arguta*). The assembled genome of kiwifruit contains 39,040 genes and has a total length of 616.1 Mb (Huang *et al*., 2013). Numerous domesticated species exist in *Actinidia* that are economically important, including *A. deliciosa, A. chinensis, A. eriantha* and *A. arguta* (Atkinson and Macrae, 2007). However, the present globally cultivated kiwifruit is limited to *A. deliciosa* and *A. chinensis*, thence the genetic background of the cultivated kiwifruits is limited (Oh *et al*., 2019). The commercially cultivated area of the crop worldwide is around 4 lakh ha with an annual production of about 6 million tonnes (FAOSTAT, 2018). *Actinidia* species have ample dietary fibre, vitamin C and various minerals in its fruit (Park *et al*., 2013). Based on the flower types, kiwifruit genotypes are categorised into two groups, i.e. staminate (male) and pistillate (female). Staminate genotypes are Tomuri and Allision & pistillate are Monty, Hayward, Bruno, Allision and Abbott.

Genetic diversity of crops is the raw material for breeding new crop varieties in response to the requirements of diverse agricultural systems (Brussaard *et al*., 2010). Assessing the pattern and level of the genetic differences for conserved genetic resources or for crop cultivars is thus crucial, in particular, for assisting the assortment of parental combinations to create hybrids with superior agronomic traits (Glaszmann *et al*., 2010) and developing conservation stratagems to obstruct genetic erosion during breeding and domestication programs (Frankham, 2010; Gepts, 2006). Systematic management of genetic resources for utilization or preservation in plant breeding programs needs fast and accurate analysis of genetic diversity levels and degree of genetic relatedness. Phenotypic distinction based on the explication of the physiological and morphological traits can be used, but this method necessitates the considerable observation of plants till maturity (Palombi and Damiano, 2002). Hence, genetic diversity is rather assessed by DNA analysis techniques and for this, numerous DNA markers are used. Polymerase Chain Reaction (PCR) based random amplified polymorphic DNA (RAPD) markers have been tremendously used for DNA fingerprinting in plants. These markers are mostly dominant and detect variation in both coding and non-coding region of the genome (Sedra *et al*., 1998). RAPD analysis is technically simple and provides an approach to characterize different genotypes.

Also, the identification of male and female genotypes and of diverse cultivars within the species is the initial phase towards the precise classification of kiwifruit germplasm. Several breeding programmes have been introduced to develop novel cultivars. However, dioecy represents an important restraint. Kiwi plants can be propagated by seeds or from vegetative parts. To maintain genetic purity and uniformity of the cultivars, these are conventionally propagated through rootstock on which commercial cultivars can be grafted or budded. The kiwifruit was preliminary raised from seed, which resulted in the progeny of highly variable plants but the vital issue is the need to identify the pistillate and staminate cultivars until they flower which takes 3-4 years. Due to this, if seeds are used for propagation, then farmers are not able to adequately cultivate a large number of productive female plants with only a minimal number of male plants. Identification of sex in the early stage is a big problem in kiwifruit. Sex-linked markers can reduce the labour, time, and costs associated with breeding programmes, and facilitate examining the sex-determination systems. Sex-linked molecular markers are useful in breeding programs and allow the understanding of dioecism in kiwifruit. Also, the molecular markers linked to the specific genotypes can help in their easy identification.

So, the present study was carried out to study the genetic diversity among kiwifruit genotypes and to find male and female-specific RAPD markers to ease the identification of the sex of the plants at the juvenile stage so that the material can be raised as male and female populations separately.

## MATERIAL AND METHODS

### Plant Material

Kiwifruit is a relatively new establishment in India and so far, there are only seven commercially grown cultivars. All these cultivars are being perpetuated at the experimental orchard of the Department of Pomology, Dr. Y.S. Parmar University of Horticulture and Forestry, Nauni-Solan and they are used as the experimental material. Among these, two are staminate viz. ‘Tomuri’ and ‘Allision’ and five are pistillate viz. ‘Hayward’, ‘Bruno’, ‘Monty’, ‘Allision’ and ‘Abbott’. Young green leaves were collected from the different vines of each cultivar and genomic DNA was isolated from them.

### Genomic DNA isolation, purification and quantification

Genomic DNA was isolated from the leaf samples of seven kiwifruit genotypes using CTAB extraction method of Saghai-Maroof *et al*. (1984) with little modifications. The DNA was isolated from each of the samples and five samples of each genotype from different vines were pooled together for further steps. Extracted DNA was purified to remove RNA, polysaccharides, phenols and proteins by giving RNase, phenol: chloroform: isoamyl alcohol (25: 24: 1) and proteinase K treatments.

The DNA isolated from each genotype was run on 0.8% agarose gel and analysed using nanodrop (Thermo Scientific) and finally, the dilution of the isolated DNA was done up to 80ng/µl and subjected to polymerase chain reaction (PCR) amplification.

### PCR conditions

A total of 94 decamer primers of random sequences (RAPD primers) were obtained from Eurofins Genomics to study genetic diversity among the kiwifruit genotypes. Amplification was carried out in 20 µl reaction mixture containing 80 ng genomic DNA, 3 mM MgCl^2^, 10 X PCR buffer, 0.25 mM dNTPs, 0.5 µM primer and 0.05 U Taq DNA Polymerase. Benchtop™ Lab Systems (programmable thermal cycler-BT-B960) was used for PCR amplification which consisted of initial denaturation at 94° for 5 min followed by 35 cycles of denaturation at 94° for 1 min, annealing at 26° C to 32° C (depending on the primer) for 1 min, extension at 72° C for 1 min and final extension at 72° C for 5 min. The amplified DNA fragments were resolved on ethidium bromide stained 2.0% (w/v) agarose gel and the amplified products were visualised using ‘Benchtop Lab Systems Gel Documentation System’.

### Allele scoring and Data Analysis

The amplified bands were scored visually on the basis of presence and absence of bands (taken as 1 and 0 respectively) for each kiwifruit genotype. The amplified band sizes were determined on the basis of their migration with respect to the marker (100 bp DNA ladder) by using ‘Fusion Solo’ software (Vilber Lourmat, France). ‘SIMQUAL’ sub-program of the software NTSYS-pc (Numerical Taxonomy and Multivariate Analysis System) was used to calculate the similarity genetic distance and cluster analysis was done to evaluate the relationship among the different kiwifruit genotypes. The UPGMA (Unweighted pair-group method with Arithmetic Mean) sub-program of NTSYS-pc was used to construct dendrogram using distance matrix and ‘EIGEN’ sub-program was used for Principle Component Analysis (PCA) and to construct both 2D and 3D PCA plots.

## RESULTS AND DISCUSSION

A total of 94 primers were screened for amplifying the genomic DNA of the kiwifruit genotypes out of which 23 showed amplification in all the kiwifruit genotypes (Table 1).

**Table 1:**
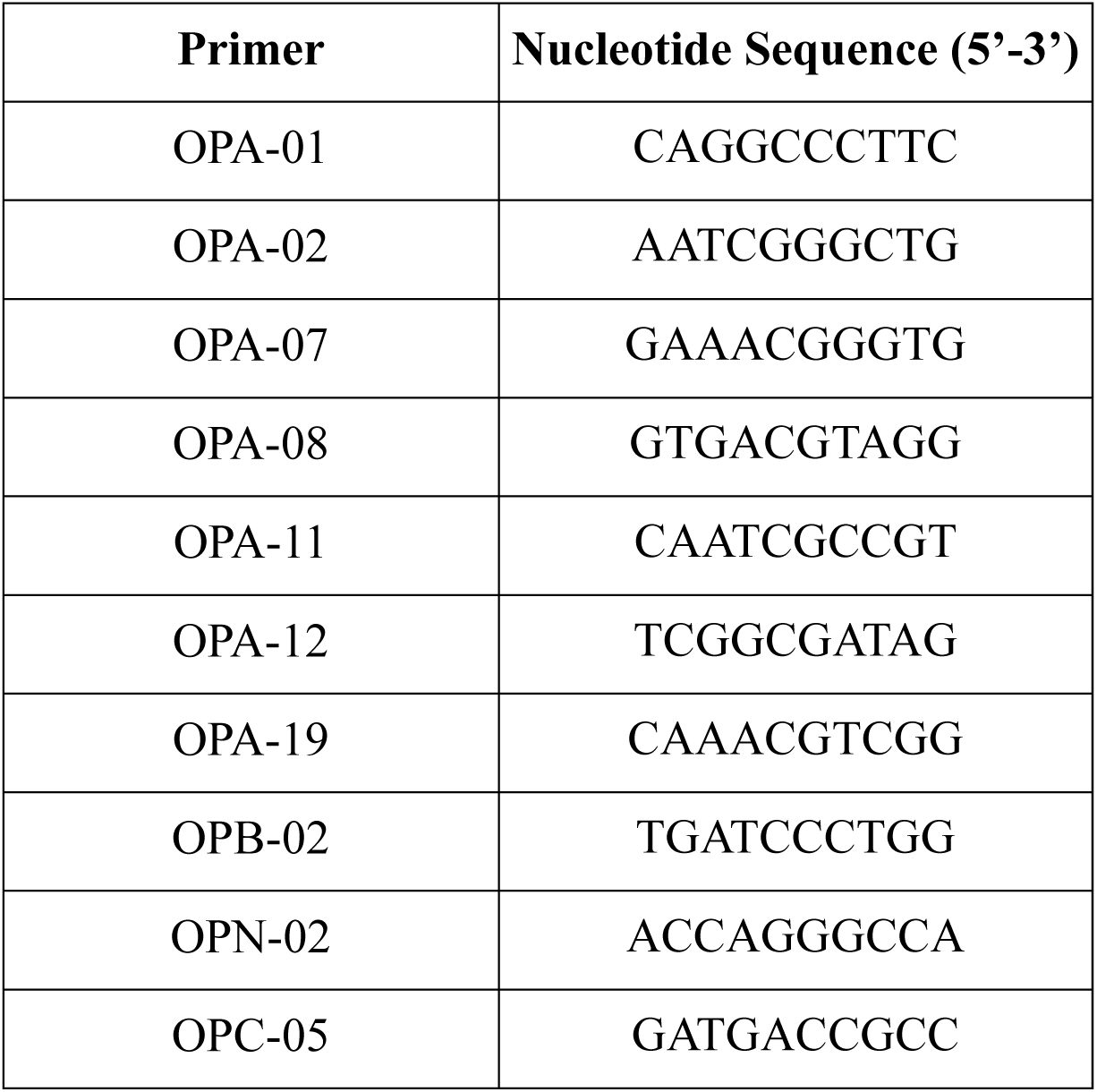
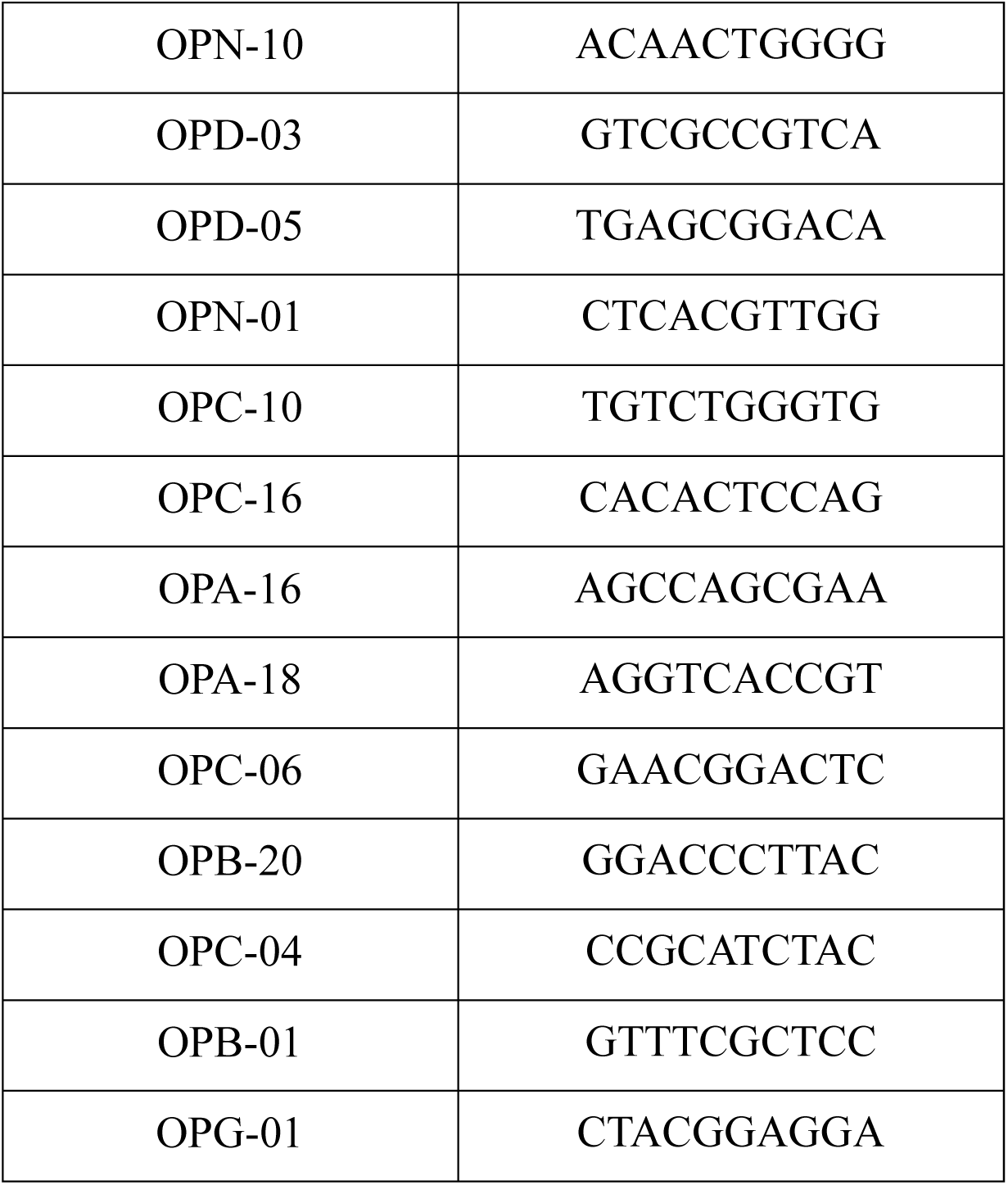
Nucleotide sequences of the primers which showed amplification among all the kiwifruit genotypes

These primers were further used for generating the DNA fingerprint profile of kiwifruit genotypes and their Polymorphism Information Content (PIC) values were calculated (Table 2). The PIC values ranged from 0.91 to 0.95 indicating the high informativeness of these primers for the assessment of genetic diversity in kiwifruit genotypes, as the PIC value higher than 0.7 is highly informative while a value of 0.44 is considered to be fairly informative (Hildebrand *et al*., 1994).

**Table 2:** Genomic DNA amplification products and the PIC values generated in kiwifruit genotypes

| **Primer** | **No. of Alleles** | **Amplicon Size Range (bp)** | **PIC** |
| --- | --- | --- | --- |
| OPA-01 | 17 | 100-1200 | 0.933 |
| OPA-02 | 19 | 200-1200 | 0.930 |
| OPA-07 | 24 | 300-1200 | 0.944 |
| OPA-08 | 23 | 300-1200 | 0.932 |
| OPA-11 | 17 | 300-1200 | 0.925 |
| OPA-12 | 20 | 300-1200 | 0.937 |
| OPA-19 | 27 | 200-1200 | 0.952 |
| OPB-02 | 18 | 300-1200 | 0.932 |
| OPN-02 | 15 | 400-1100 | 0.922 |
| OPC-05 | 15 | 300-1100 | 0.921 |
| OPN-10 | 18 | 300-1200 | 0.936 |
| OPD-03 | 16 | 200-1100 | 0.924 |
| OPD-05 | 25 | 200-1200 | 0.946 |
| OPN-01 | 17 | 200-1100 | 0.928 |
| OPC-10 | 16 | 100-1200 | 0.923 |
| OPC-16 | 26 | 200-1200 | 0.956 |
| OPA-16 | 13 | 300-900 | 0.921 |
| OPA-18 | 19 | 400-1100 | 0.914 |
| OPC-06 | 19 | 300-1200 | 0.937 |
| OPB-20 | 20 | 200-1200 | 0.942 |
| OPC-04 | 15 | 100-1100 | 0.933 |
| OPB-01 | 24 | 200-1200 | 0.914 |
| OPG-01 | 19 | 200-1100 | 0.926 |

### Similarity Matrices

The similarity genetic distance among the kiwifruit genotypes revealed their genetic relatedness (Table 3). Tomuri (M) and Allision (M) showed the maximum similarity, while the most diverse genotypes were found to be Allision (M) and Abbott (F). Being the most diverse, Allision (M) and Abbott (F) can generate the most heterotic combination in the hybridisation program.

**Table 3:** Similarity matrix data of kiwifruit genotypes

|  | **Tomuri(M)** | **Bruno(F)** | **Allision(F)** | **Abbott(F)** | **Hayward(F)** | **Monty(F)** | **Allision(M)** |
| --- | --- | --- | --- | --- | --- | --- | --- |
| **Tomuri(M)** | 1 |  |  |  |  |  |  |
| **Bruno(F)** | 0.585 | 1 |  |  |  |  |  |
| **Allision(F)** | 0.582 | 0.662 | 1 |  |  |  |  |
| **Abbott(F)** | 0.538 | 0.574 | 0.637 | 1 |  |  |  |
| **Hayward(F)** | 0.596 | 0.671 | 0.656 | 0.629 | 1 |  |  |
| **Monty(F)** | 0.588 | 0.664 | 0.667 | 0.656 | 0.681 | 1 |  |
| **Allision(M)** | **0.719** | 0.568 | 0.571 | **0.521** | 0.607 | 0.640 | 1 |

### Cluster Tree Analysis

The phylogenetic tree was constructed to illustrate the relationship among the kiwifruit genotypes by using their similarity genetic distance data. The cluster tree analysis illustrated similar observations as shown by the similarity matrix data, which are represented in the form of a dendrogram (Fig. 1). There were two major clusters at 0.58 value of similarity coefficient. Cluster I had all the male genotypes whereas all the female genotypes fell in cluster II. These results imply that the male and female kiwifruit genotypes are diverse genetically. Shirkot *et al*. (2002) also described that cluster analysis of kiwifruit genotypes by UPGMA method exhibited two main clusters. Cluster I with two male genotypes (Tomuri and Allision) and cluster II with all the female genotypes (Hayward, Allision, Monty, Bruno and Abbott).

**Fig. 1:**
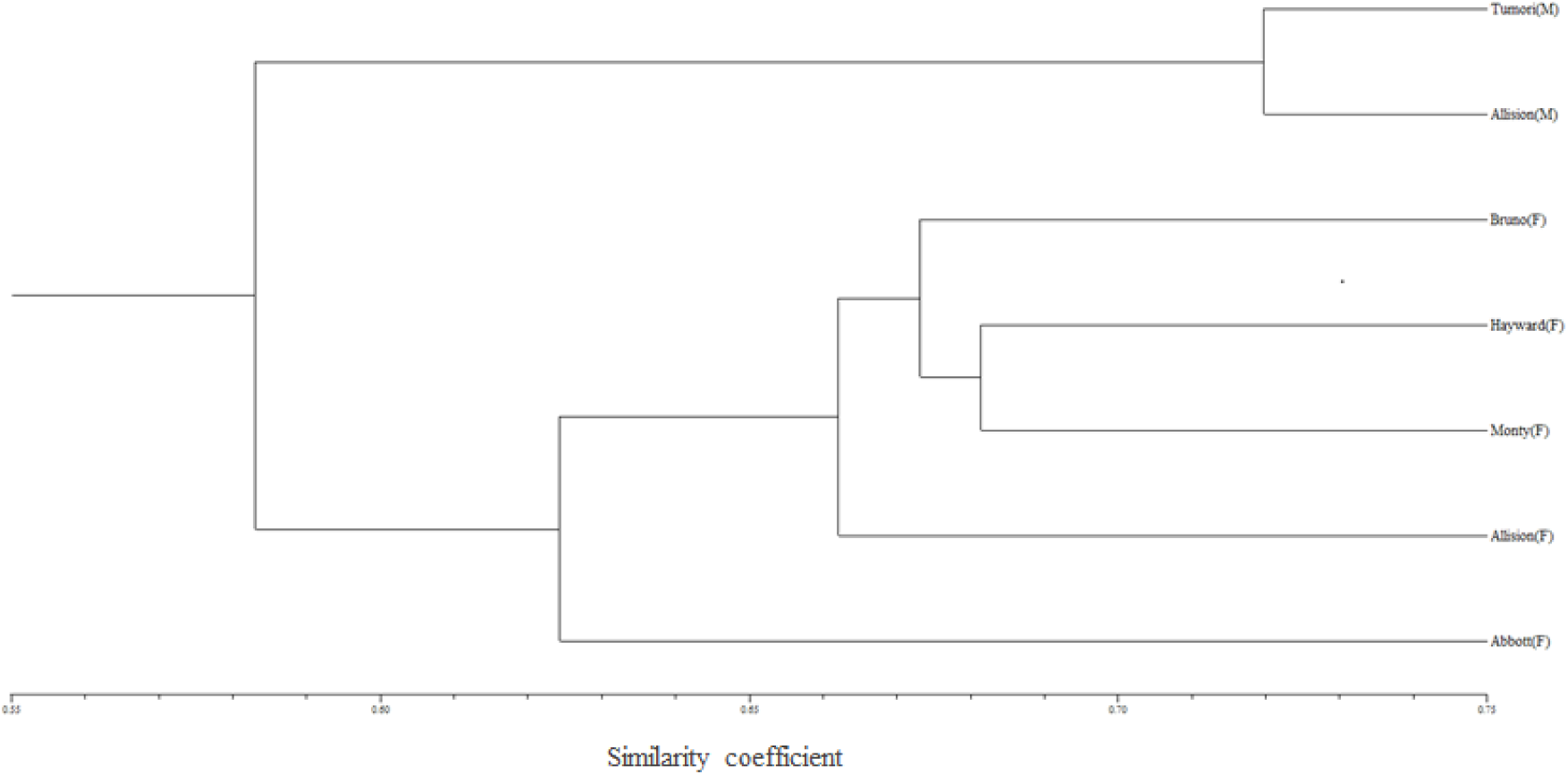
Genetic relatedness among the kiwifruit genotypes based on RAPD analysis.

### 2D and 3D Principle Component Analysis

The 2D and 3D PCA plots obtained from PCA analysis also showed similar clustering (Fig. 2 and 3), thus confirming the results of the cluster tree analysis.

**Fig. 2:**
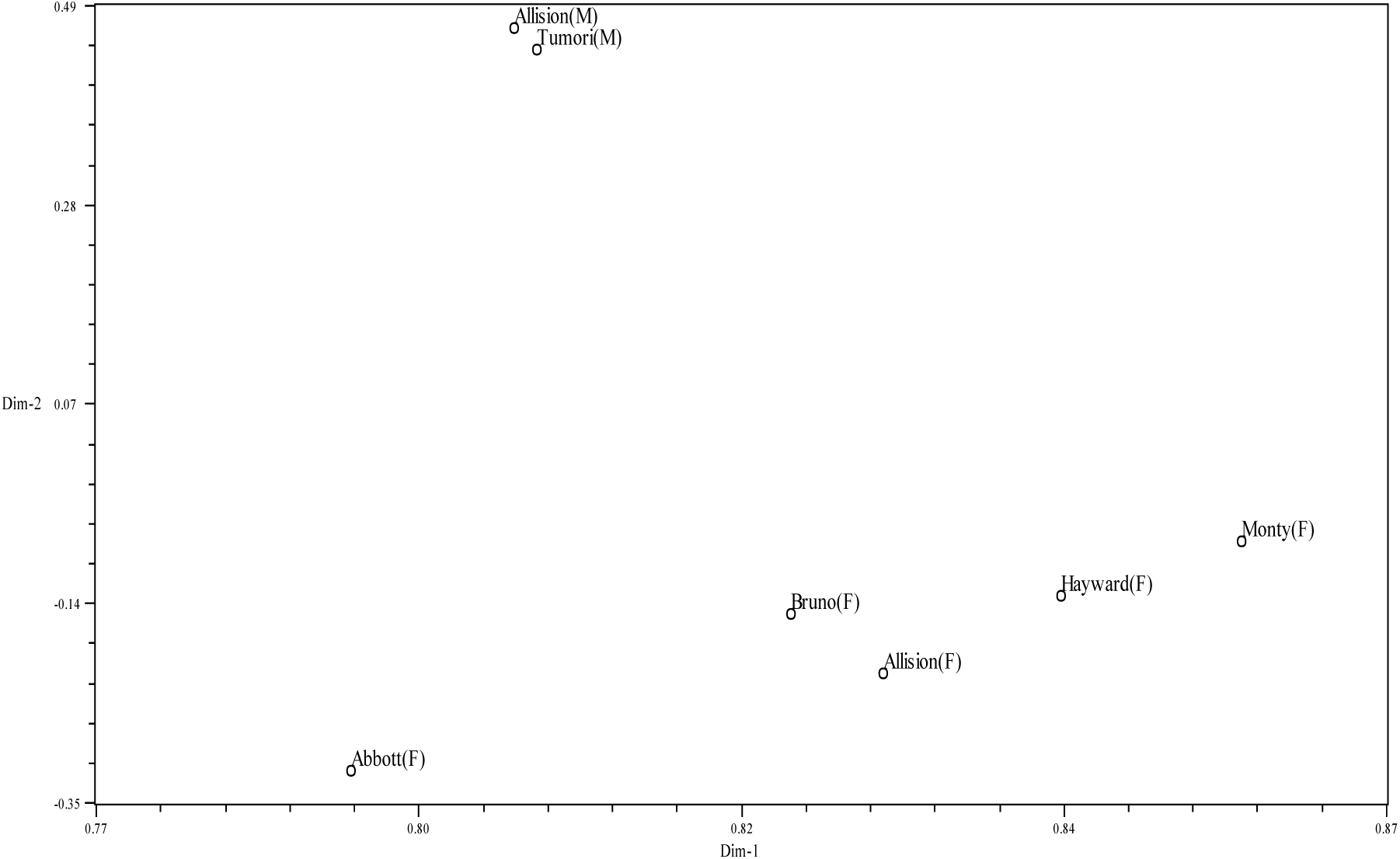
2D PCA plot of kiwifruit genotypes.

**Fig. 3:**
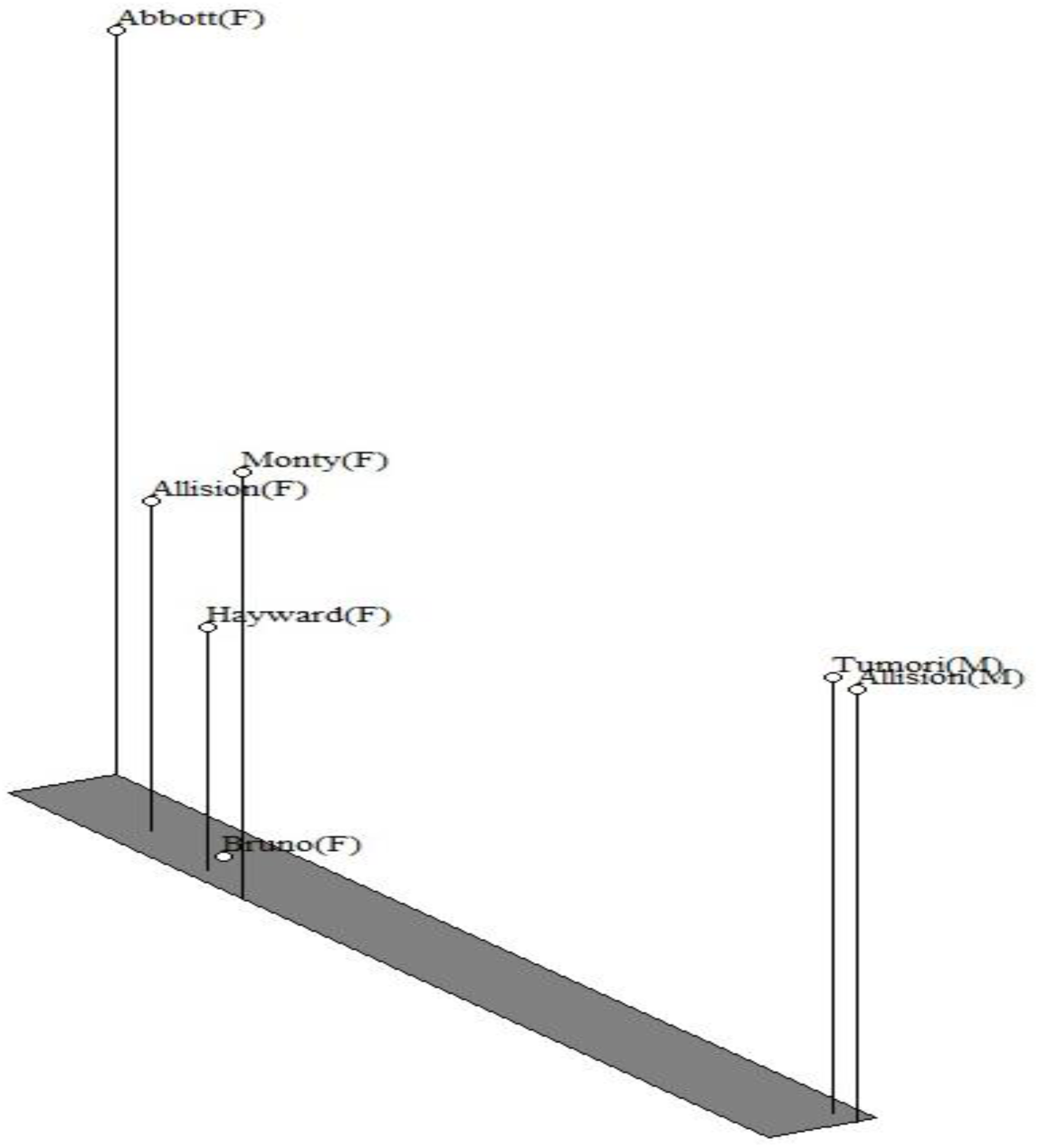
3D PCA plot of kiwifruit genotypes.

Out of the 23 primers showing amplification in all the genotypes, six primers showed unique banding patterns. Primers OPA-01, OPA-02, OPN-02, OPC-05, OPB-02, and OPB-20 amplified unique amplicons in specific genotypes (Table 4). From these six primers, two primers can be used to differentiate the male and female kiwifruit genotypes (Table 5) viz. OPA-02 and OPN-02 (Plate 1) and four primers can be used to distinguish the specific kiwifruit genotypes (Table 6) viz. OPA-01, OPC-05 (Plate 2), OPB-02 and OPB-20. Shirkot *et al*., (2002) also found two primers (OPN-01 and OPC-05) amplifying unique amplicons in the male genotypes and six primers (OPA-01, OPA-02, OPA-11, OPA-08, OPB-01 and OPA-16) amplifying unique amplicons in the female genotypes.

**Table 4:** Primers amplifying unique amplicons in specific genotypes

| Primer | Unique amplicon size (bp) | Genotype |
| --- | --- | --- |
| OPA-01 | 167 | Bruno |
|  | 300 | Hayward |
| OPA-02 | 273 | Tomuri |
|  | 330 | Allision (M) |
| OPN-02 | 378 | Allision (M) |
|  | 805 | Allision (F) |
| OPC-05 | 193 | Allision (M) |
|  | 273 | Abbott |
| OPB-02 | 425 | Abbott |
| OPB-20 | 412 | Monty |
|  | 504 | Bruno |

**Table 5:** Primers capable of distinguishing the male and female genotypes

| Primer | Amplicon size (bp) | Genotype |
| --- | --- | --- |
| OPA-02 | 273 | Tomuri (M) |
|  | 330 | Allision (M) |
| OPN-02 | 805 | Allision (F) |
|  | 378 | Allision (M) |

**Plate 1:**
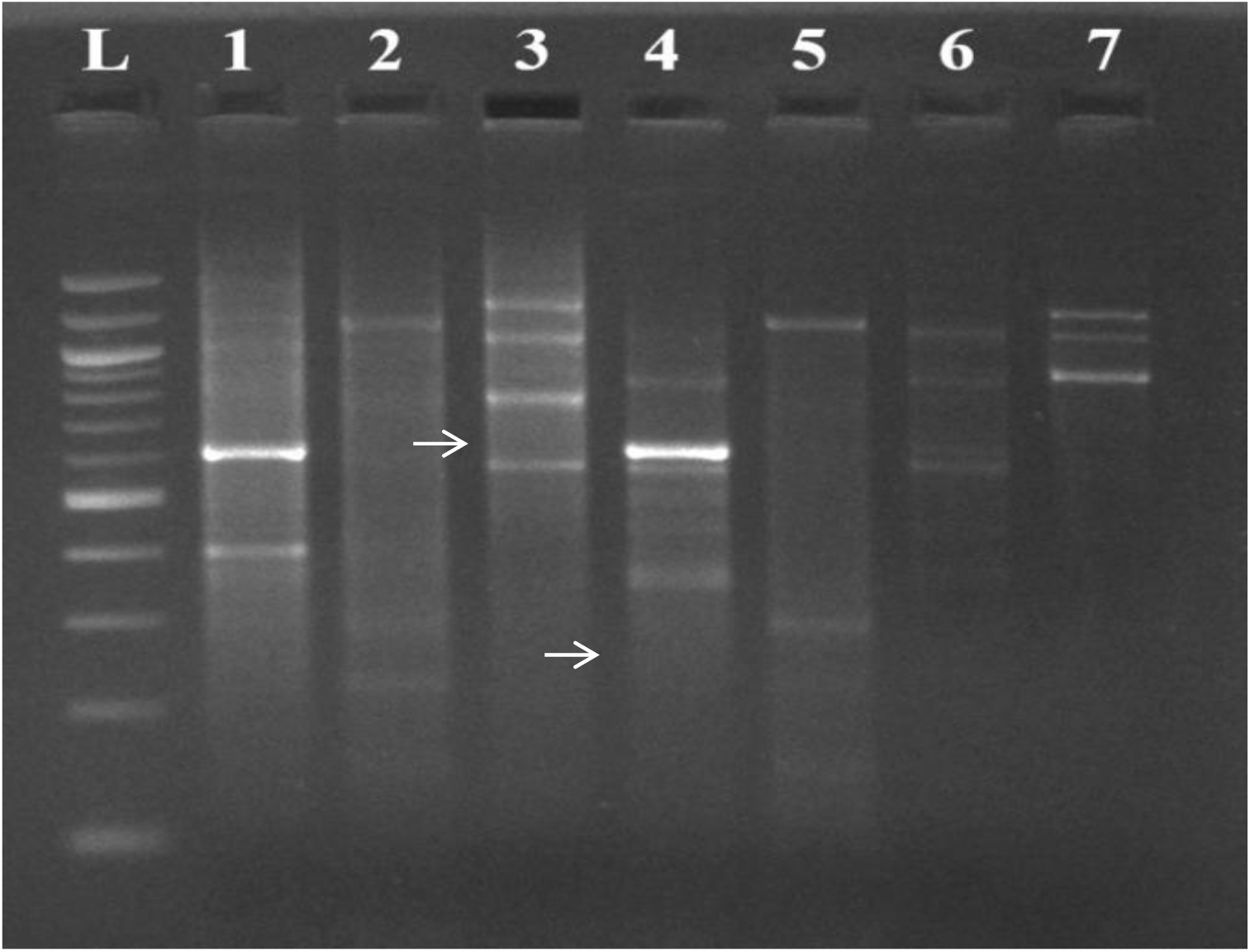
RAPD banding profile of primer OPN-02, L: 100bp DNA marker, Lanes 1: Tomuri, Lane 2: Bruno, Lane 3: Allision(F), Lane 4: Allision(M), Lane 5: Hayward, Lane 6: Monty and Lane 7: Abbott (Arrows showing **805** bp and **378** bp unique amplicons in lane 3 and lane 4 respectively)

**Plate 2:**
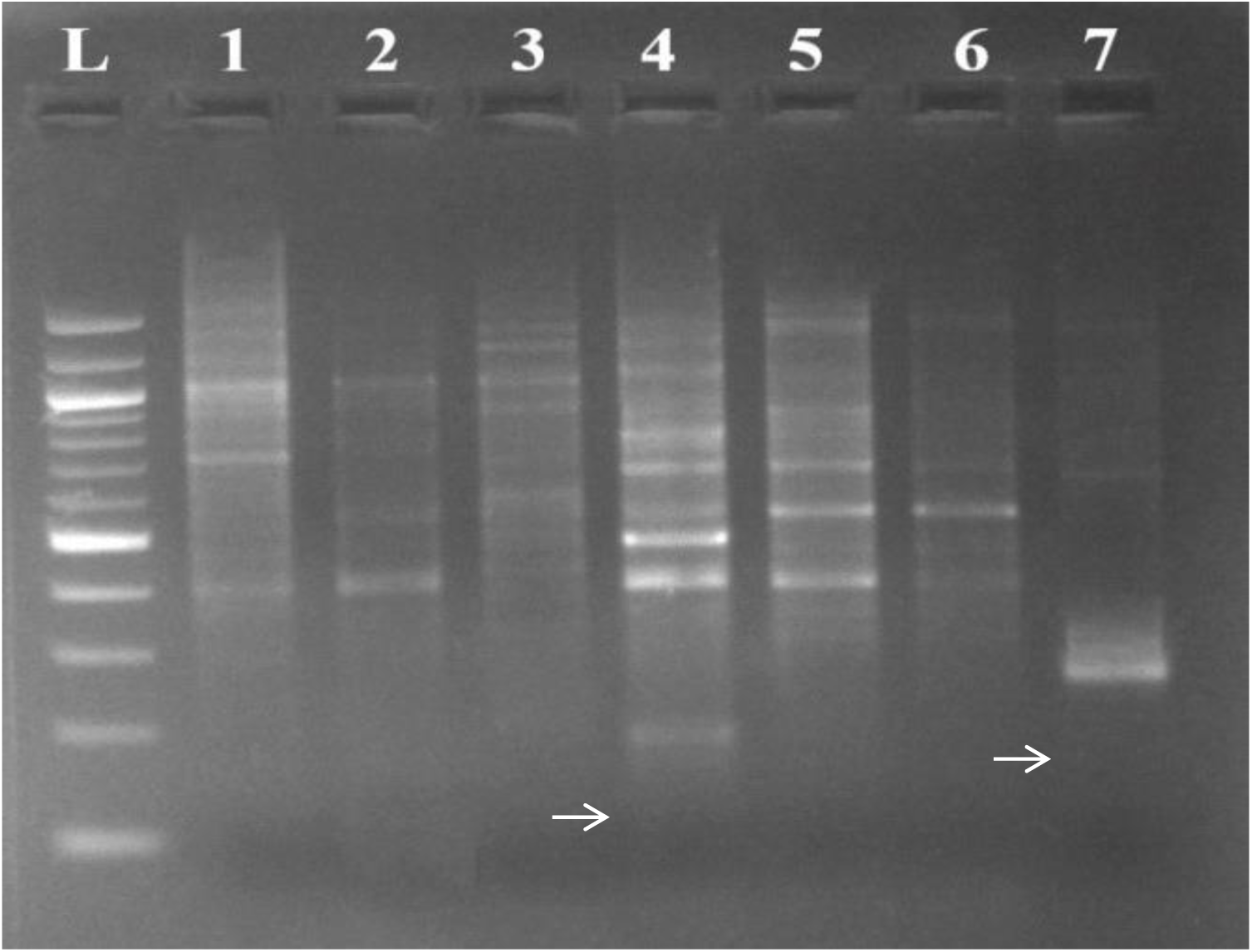
RAPD banding profile of primer OPC-05, L: 100bp DNA marker, Lanes 1: Tomuri, Lane 2: Bruno, Lane 3: Allision(F), Lane 4: Allision(M), Lane 5: Hayward, Lane 6: Monty and Lane 7: Abbott (Arrows showing **193** bp and **273** bp unique amplicons in lane 4 and lane 7 respectively)

## CONCLUSION

The generated DNA profile of the kiwifruit revealed that the primers OPA-02 and OPN-02 may be validated as putative sex-linked markers as these have the potential to be further used for the characterization of the male and female kiwifruit genotypes. Also, the variations found in the closely related genotypes showed the efficiency of the RAPDs for diversity analysis and genotyping of kiwifruit genotypes over the morphological and isozyme markers.

## Author Approvals

All authors have seen and approved the manuscript and has not been accepted or published elsewhere.

## REFERENCES

Ferguson, A. R. (1991). Kiwifruit (*Actinidia*). Genetic Resources of Temperate Fruit and Nut Crops, 290: 603–656.

Yan, G., Yao, J., Ferguson, A. R., McNeilage, M. A., Seal, A. G., & Murray, B. G. (1997). New reports of chromosome numbers in *Actinidia* (Actinidiaceae). New Zealand Journal of Botany, 35(2): 181–186.

Huang, S., Ding, J., Deng, D., Tang, W., Sun, H., Liu, D & Yu, J. (2013). Draft genome of the kiwifruit *Actinidia chinensis*. Nature communications, 4: 2640–2649.

FAOSTAT. (2018). Available online: http://www.fao.org/faostat/en/#data.

Palombi, M. and Damiano, C. (2002). Comparison between RAPD and SSR molecular markers in detecting genetic variation in kiwifruit (*Actinidia deliciosa* A. Chev.). Plant Cell Reports, 20(11): 1061–1066.

Sedra, M.H., Lashermes, P., Trouslot, P. and Combes, M.C. (1998). Identification and genetic diversity analysis of date palm (*Phoenix dactylifera* L.) varieties from Morocco using RAPD markers. Euphytica, 103(1): 75–82.

Saghai-Maroof, M. A., Soliman, K. M., Jorgensen, R. A. and Allard, R. W. (1984). Ribosomal DNA spacer-length polymorphisms in barley: Mendelian inheritance, chromosomal location, and population dynamics. Proceedings of the National Academy of Sciences, 81(24): 8014–8018.

Hildebrand, C. E., David, C., Torney, C. & Wagner, P. (1994). Informativeness of polymorphic DNA markers. Los Alamos Sci, 20: 100–102.

Shirkot, P., Sharma, D. R. & Mohapatra, T. (2002). Molecular identification of sex in *Actinidia deliciosa* var. deliciosa by RAPD markers. Scientia Horticulturae, 94(2): 33–39.

Atkinson, R. G., & MacRae, E. A. (2007). I. 13 Kiwifruit. Biotechnology in agriculture and forestry, 60, 329–346.

Oh, S., Lee, M., Kim, K., Han, H., Won, K., Kwack, Y. B., … & Kim, D. (2019). Genetic diversity of kiwifruit (Actinidia spp.), including Korean native A. arguta, using single nucleotide polymorphisms derived from genotyping-by-sequencing. Horticulture, Environment, and Biotechnology, 60(1); 105–114.

Park, Y. S., Im, M. H., Ham, K. S., Kang, S. G., Park, Y. K., Namiesnik, J., … & Gorinstein, S. (2013). Nutritional and pharmaceutical properties of bioactive compounds in organic and conventional growing kiwifruit. Plant Foods for Human Nutrition, 68(1); 57–64.

Brussaard, L., Caron, P., Campbell, B., Lipper, L., Mainka, S., Rabbinge, R., … & Pulleman, M. (2010). Reconciling biodiversity conservation and food security: scientific challenges for a new agriculture. Current opinion in Environmental sustainability, 2(2); 34–42.

Glaszmann, J. C., Kilian, B., Upadhyaya, H. D., & Varshney, R. K. (2010). Accessing genetic diversity for crop improvement. Current opinion in plant biology, 13(2); 167–173.

Frankham, R. (2010). Challenges and opportunities of genetic approaches to biological conservation. Biological conservation, 143(9); 1919–1927.

Gepts, P. (2006). Plant genetic resources conservation and utilization: the accomplishments and future of a societal insurance policy. Crop science, 46(5); 2278–2292.

